# Differential Protection against Acetaminophen-induced Liver and Kidney Injury by N-Acetylcysteine and Fomepizole in Female Mice

**DOI:** 10.64898/2026.10.16.759774

**Authors:** Yasaman Etemadi, Timothy A. Fields, Anup Ramachandran, Hartmut Jaeschke

**Affiliations:** Department of Pharmacology, Toxicology, and Therapeutics, and; Department of Pathology and Laboratory Medicine, University of Kansas Medical Center, Kansas City, KS, 66160, USA

**Author notes:** For Correspondence: Hartmut Jaeschke, PhD, Department of Pharmacology, Toxicology, and Therapeutics University of Kansas Medical Center, 3901 Rainbow Blvd, MS 1018 Kansas City, KS, 66160, USA.

**Keywords:** Acute kidney injury, acute liver failure, N-acetylcysteine, buthionine sulfoximine, glutathione, 4-methylpyrazole, Fomepizole

## Abstract

Acetaminophen (APAP) overdose is the leading cause of acute liver failure and is frequently complicated by acute kidney injury (AKI), which worsens clinical outcomes. Although N-acetylcysteine (NAC) effectively attenuates APAP-induced hepatotoxicity by restoring hepatic glutathione (GSH), its efficacy against AKI remains limited, suggesting distinct mechanisms of renal injury. We investigated the role of GSH homeostasis and potential beneficial effects of NAC and Fomepizole (4-methylpyrazole) in APAP-induced liver and kidney injury using female C57BL/6J mice, which naturally lack renal CYP2E1 expression and therefore provide a unique model to examine nephrotoxicity independent of renal CYP2E1-mediated bioactivation. Female mice received APAP (300 or 600 mg/kg) with or without buthionine sulfoximine (BSO, 50 mg/kg) to inhibit GSH synthesis, NAC (500 mg/kg) to promote GSH recovery, or Fomepizole (50 mg/kg) to inhibit CYP2E1-mediated bioactivation. APAP overdose caused severe hepatotoxicity, whereas kidney injury remained minimal during the first 24 h. Although BSO markedly enhanced hepatic GSH depletion and liver injury, it did not significantly worsen renal dysfunction or tubular injury. In contrast, NAC substantially reduced liver injury and improved survival after severe APAP overdose; yet renal dysfunction progressed by 48 h, accompanied by tubular vacuolation, increased KIM-1 and osteopontin expression, and tubular cell death. Fomepizole provided substantial protection against injury in both organs. These findings demonstrate that delayed APAP-induced AKI can develop independent of the severity of liver injury, even with effective NAC-mediated hepatoprotection. On the other hand, Fomepizole protects against AKI in female mice by a CYP2E1-independent mechanism.

## 1. Introduction

Acetaminophen (APAP) is one of the most widely used analgesic and antipyretic drugs worldwide. However, APAP overdose remains the leading cause of ALF in the United States and many Western countries (Lee, 2004). In addition to hepatotoxicity, APAP overdose is frequently complicated by acute kidney injury (AKI), which develops in approximately 30-50% of patients and substantially increases morbidity and one-year mortality following ALF (Antoine et al., 2015; Chen et al., 2015; Hadem et al., 2019; Mazer & Perrone, 2008; Stollings et al., 2016; Tujios et al., 2015). Clinical studies have shown that patients who develop AKI exhibit elevated blood urea nitrogen and serum creatinine levels and have significantly poorer one-year survival than patients without renal dysfunction (Antoine et al., 2015; Hadem et al., 2019; Tujios et al., 2015).

Consistent with these observations, our previous clinical analysis demonstrated that APAP overdose patients who developed renal failure also exhibited more severe hepatocellular injury than those without AKI (Akakpo et al., 2020). Although N-acetylcysteine (NAC) effectively protects against APAP-induced hepatotoxicity by restoring glutathione (Corcoran & Wong, 1986), its efficacy in preventing AKI remains limited, suggesting that the mechanisms responsible for kidney injury differ from those driving liver injury (Davenport & Finn, 1988; Hengy et al., 2009; Jones & Vale, 1993; Slitt et al., 2004). To further define the relationship between liver and kidney injury following APAP overdose, our previous experimental study in male mice demonstrated that pharmacological depletion of glutathione using buthionine sulfoximine (BSO) markedly exacerbated APAP-induced kidney injury without significantly increasing hepatotoxicity, whereas NAC substantially reduced liver injury but failed to prevent AKI (Etemadi et al., 2026a). These findings indicated that liver and kidney injury can be differentially regulated following APAP overdose. However, because those studies were performed in male mice expressing CYP2E1 in both the liver and kidney, they could not determine whether renal CYP2E1-dependent bioactivation is required for the development of AKI. The mechanisms underlying APAP-induced liver injury have been extensively characterized and are initiated by CYP2E1-mediated metabolic activation of APAP (McGill and Jaeschke, 2013). During overdose, excessive formation of the reactive metabolite N-acetyl-p-benzoquinone imine (NAPQI) rapidly depletes hepatic GSH and covalently binds to mitochondrial proteins, triggering mitochondrial dysfunction, oxidant stress, and extensive hepatocellular necrosis (Gujral et al., 2002; Jaeschke & Ramachandran, 2024; Jaeschke et al., 2019). In contrast, the mechanisms responsible for APAP-induced AKI remain considerably less well defined. Although CYP2E1 expressed in proximal tubular cells contributes to local APAP bioactivation and NAPQI formation (Akakpo et al., 2020, 2023; Hart et al., 1994, 1995), accumulating evidence indicates that renal injury differs fundamentally from hepatotoxicity. Rather than mitochondrial dysfunction, APAP nephrotoxicity is characterized primarily by endoplasmic reticulum (ER) stress and apoptotic signaling (Akakpo et al., 2023; Lorz et al., 2004). Pharmacological inhibition of caspases with Ac-DEVD-CHO markedly attenuates kidney injury without affecting liver injury, highlighting fundamental differences between the mechanisms governing APAP-induced liver and kidney injury (Akakpo et al., 2023). Female C57BL/6J mice provide a unique physiological model to address this unresolved question because they naturally lack CYP2E1 expression in proximal tubular cells due to the absence of testosterone-dependent induction (Hoivik et al., 1995; Hu et al., 1993; Pan et al., 1992). Consequently, renal APAP bioactivation is largely absent while hepatic CYP2E1 expression remains intact, allowing direct evaluation of whether APAP-induced AKI requires local renal CYP2E1-dependent bioactivation. Therefore, the objective of this study was to determine whether modulation of glutathione homeostasis with BSO or NAC alters APAP-induced kidney injury in female mice lacking renal CYP2E1, and whether CYP2E1 inhibition with Fomepizole (4-methylpyrazole; 4-MP) protects against AKI in the absence of renal bioactivation.

## 2. Materials and methods

### 2.1. Animals

Eight- to ten-week-old female C57BL/6J mice were purchased from Jackson Laboratory (Bar Harbor, ME, USA) and maintained under standard housing conditions with unrestricted access to food and water. Animals were housed in a temperature-controlled facility on a 12-hour light/dark cycle. All experimental procedures were approved by the Institutional Animal Care and Use Committee (IACUC) at the University of Kansas Medical Center and conducted in accordance with the National Research Council’s Guide for the Care and Use of Laboratory Animals. All animal procedures reported in the study complied with the ARRIVE guidelines.

### 2.2. Experimental design

Unless otherwise specified, all reagents were obtained from Sigma-Aldrich (St. Louis, MO, USA). Mice were fasted overnight for 16 h before treatment. APAP was dissolved in warm saline and administered intraperitoneally (i.p.) at doses of 300 or 600 mg/kg. For glutathione depletion studies, mice received buthionine sulfoximine (BSO; 50 mg/kg, i.p.) 2 h before APAP administration (300 mg/kg, i.p.). For glutathione replenishment studies, N-acetylcysteine (NAC; 500 mg/kg, i.p.) was administered concurrently with APAP (600 mg/kg, i.p.). In another experiment, 4-methylpyrazole (4-MP; 50 mg/kg, i.p.) was administered concurrently with APAP (600 mg/kg, i.p.).

Animals were euthanized under isoflurane anesthesia at 3, 6, 24, or 48 h following APAP treatment. Blood samples were collected immediately from the inferior vena cava using heparinized syringes, and plasma was isolated by centrifugation at 18,000 × g for 2 min. Liver and kidney tissues were excised, rinsed in saline, and sectioned. Portions of each organ were fixed overnight in 10% neutral-buffered formalin for histological evaluation, while the remaining tissue was snap-frozen in liquid nitrogen and stored for subsequent molecular and biochemical analyses.

### 2.3. Biochemical assays

Plasma alanine aminotransferase (ALT) activity, a marker of hepatocellular injury, was measured using a commercially available colorimetric assay kit (MedTest, Canton, MI, USA) according to the manufacturer’s instructions. Renal function was assessed by measuring blood urea nitrogen (BUN) and creatinine concentrations using QuantiChrom™ Urea and Creatinine Assay Kits, respectively (BioAssay Systems, Hayward, CA, USA). Hepatic and renal glutathione levels were determined using a modified Tietze recycling assay as previously described (Etemadi et al., 2026a).

### 2.4. Histology and Immunohistochemistry

For histopathological evaluation, liver and kidney tissues were fixed in 10% neutral-buffered formalin, paraffin embedded and sectioned at a thickness of 5 μm. Tissue sections were stained with hematoxylin and eosin (H&E) or Periodic Acid–Schiff (PAS) using standard histological procedures to assess tissue morphology and injury. Renal tubular injury was evaluated by a renal expert pathologist (T.A.F.) using a blinded histopathological scoring system focused on the renal cortex and outer stripe of the outer medulla (OSOM), regions enriched in proximal tubules and particularly susceptible to APAP-induced nephrotoxicity (Cummings et al., 2000; Radi, 2019). Injury was graded according to the severity and extent of tubular damage using the following criteria: Score 1, mild injury affecting <10% of the cortex; Score 2, mild injury affecting >50% of the cortex; Score 3, moderate-to-severe injury affecting <10% of the cortex; and Score 4, moderate-to-severe injury affecting >50% of the cortex. We evaluated DNA fragmentation by terminal deoxynucleotidyl transferase-mediated dUTP nick-end labeling (TUNEL) using the In Situ Cell Death Detection Kit, Fluorescein (Roche Diagnostics, Indianapolis, IN, USA; Cat# 11684795910), according to the manufacturer’s instructions.

For immunohistochemical detection of kidney injury molecule-1 (KIM-1), kidney sections were blocked for 1 h at room temperature in 3% bovine serum albumin (BSA) and incubated overnight with a rabbit anti-KIM-1 primary antibody (Cell Signaling Technology, Danvers, MA, USA; Cat# 14971). We quenched endogenous peroxidase activity with 3% hydrogen peroxide, then performed three washes in Tris-buffered saline (TBS). Immunoreactivity was visualized using the VECTASTAIN® Elite ABC detection system (Vector Laboratories, Cat# PK-4001) and a 3,3′-diaminobenzidine (DAB) substrate kit (Cell Signaling Technology, Cat# 8059). KIM-1 and osteopontin immunostaining were quantified using Fiji (ImageJ) by calculating the integrated optical density (IOD) relative to the total tissue area (IOD/tissue area). For each mouse, 3-4 randomly selected, non-overlapping fields of the renal cortex were analyzed at 20× magnification. The average value from each animal was considered one biological replicate, with 3-4 mice included per experimental group.

### 2.5. RNA isolation and quantitative PCR

Total RNA was extracted from liver and kidney tissues using TRIzol™ reagent (Invitrogen, Carlsbad, CA, USA) according to the manufacturer’s instructions. Two micrograms of total RNA were reverse transcribed into complementary DNA (cDNA) using a High-Capacity cDNA Reverse Transcription Kit (Applied Biosystems, Cat# 4368814). Quantitative real-time PCR (qPCR) was performed using PowerUp™ SYBR® Green Master Mix (Thermo Fisher Scientific, Cat# A25742) and gene-specific primers obtained from Integrated DNA Technologies (Coralville, IA, USA). The expression of kidney injury markers, including neutrophil gelatinase-associated lipocalin (NGAL) and kidney injury molecule-1 (KIM-1), was quantified using the comparative Ct method.

Gene expression levels were normalized to 18S rRNA as the endogenous control, and relative changes were calculated using the 2^−ΔΔCt method.

### 2.6. Statistical analysis

All data are expressed as mean ± standard error of the mean (SEM). Statistical analyses were performed using an unpaired two-tailed Student’s t-test for comparisons between two groups. A p-value < 0.05 was considered statistically significant. Data analysis and graph generation were conducted using GraphPad Prism version 8.0.1 (GraphPad Software, San Diego, CA, USA).

## 3. Results

### 3.1. APAP induces severe liver injury but only limited kidney injury in female mice

To determine whether APAP produces dose-dependent liver and kidney injury in female C57BL/6J mice, animals were fasted overnight and administered either 300 mg/kg (moderate overdose) or 600 mg/kg (severe overdose) APAP. At 24 hours, serum ALT activity increased markedly in both APAP-treated groups, with substantially greater elevations following the 600 mg/kg dose, indicating severe dose-dependent hepatocellular injury (Figure 1A). In contrast, evidence of renal injury at 24 h was relatively limited. BUN remained unchanged after 300 mg/kg APAP but increased significantly in mice receiving 600 mg/kg APAP (Figure 1B), whereas serum creatinine did not differ among the experimental groups (Figure 1C). Total renal glutathione levels were also not significantly altered 24 h after APAP administration (Figure 1D). Despite the minimal changes in conventional markers of renal dysfunction, KIM-1 mRNA expression, a sensitive indicator of proximal tubular injury, was increased approximately three-fold in the 600 mg/kg group compared with controls, while only a modest increase was observed following 300 mg/kg APAP (Figure 1E). Consistent with the modest induction of KIM-1 mRNA levels, TUNEL staining revealed only a very limited number of positive cells in the kidneys of APAP-treated mice at 24 h, with no substantial increase compared with controls (Figure 1F), indicating minimal kidney dysfunction but no relevant cell death. Histopathological examination of liver and kidney sections further demonstrated the organ-specific effects of APAP in female mice. Consistent with the plasma ALT data, liver sections exhibited dose-dependent hepatocellular necrosis, with greater injury in mice treated with 600 mg/kg APAP compared with the 300 mg/kg group (Figure 2). In contrast, histological evaluation of kidney sections with PAS staining revealed no apparent morphological differences among control, 300 mg/kg APAP, and 600 mg/kg APAP-treated mice at 24 h (Figure 2). Renal tubular architecture remained preserved, with no evidence of tubular necrosis or other structural abnormalities. These findings indicate that, despite significant liver injury, neither the moderate nor the severe APAP dose induced detectable morphological kidney injury in female mice within 24 h of administration.

**Figure 1.**
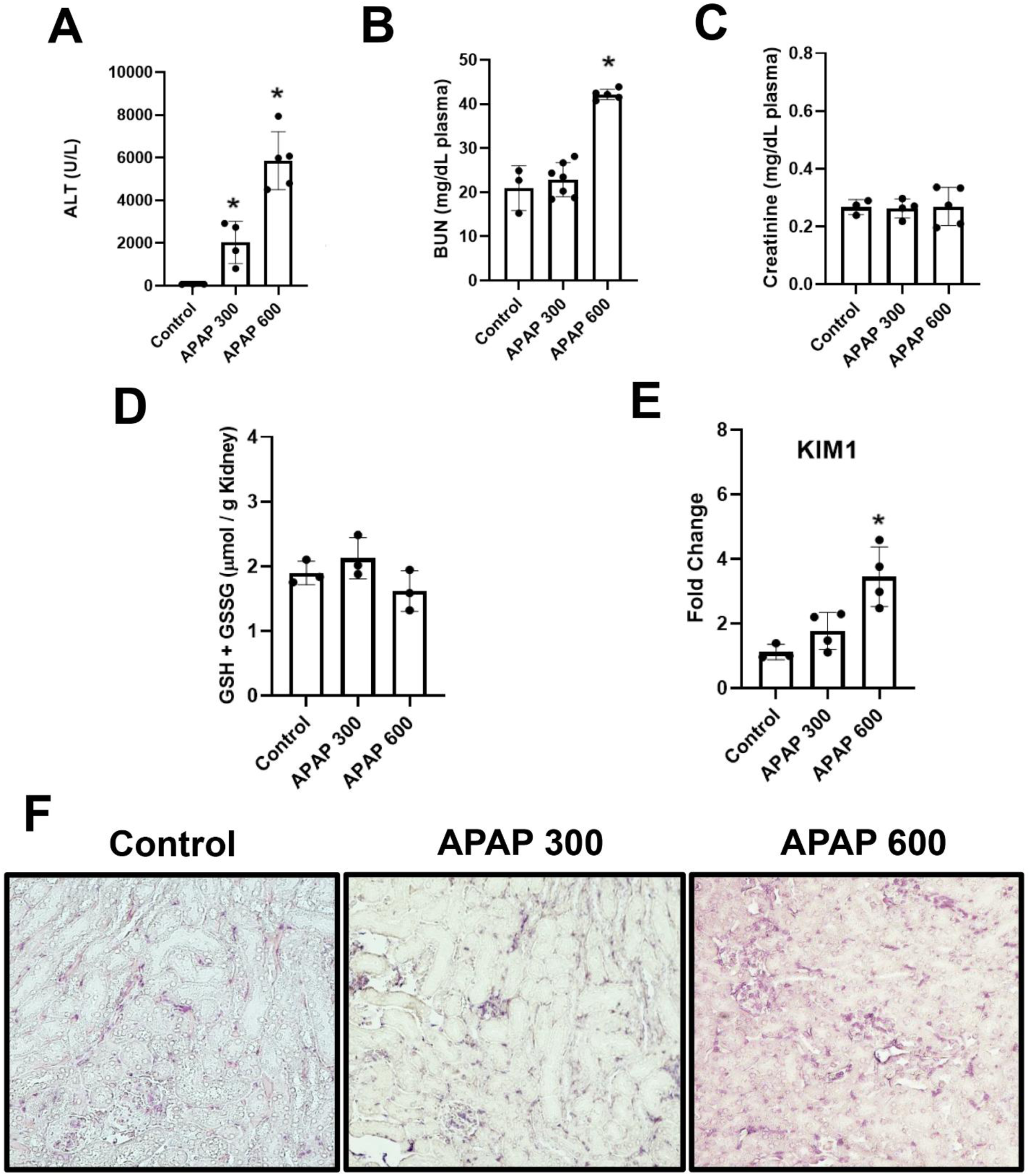
A severe APAP overdose induces marked hepatotoxicity but only limited renal dysfunction in female mice at 24. **h.** Overnight-fasted female C57BL/6J mice received a single intraperitoneal injection of acetaminophen (APAP; 300 or 600 mg/kg) and were euthanized 24 h later. (A) Plasma ALT level, (B) blood urea nitrogen (BUN), (C) plasma creatinine, (D) total renal glutathione, (E) renal Kim-1 mRNA expression, and (F) representative TUNEL-stained kidney sections (40x). Data are presented as mean ± SEM (n = 4–5 mice/group). *P* < 0.05 versus control.

**Figure 2.**
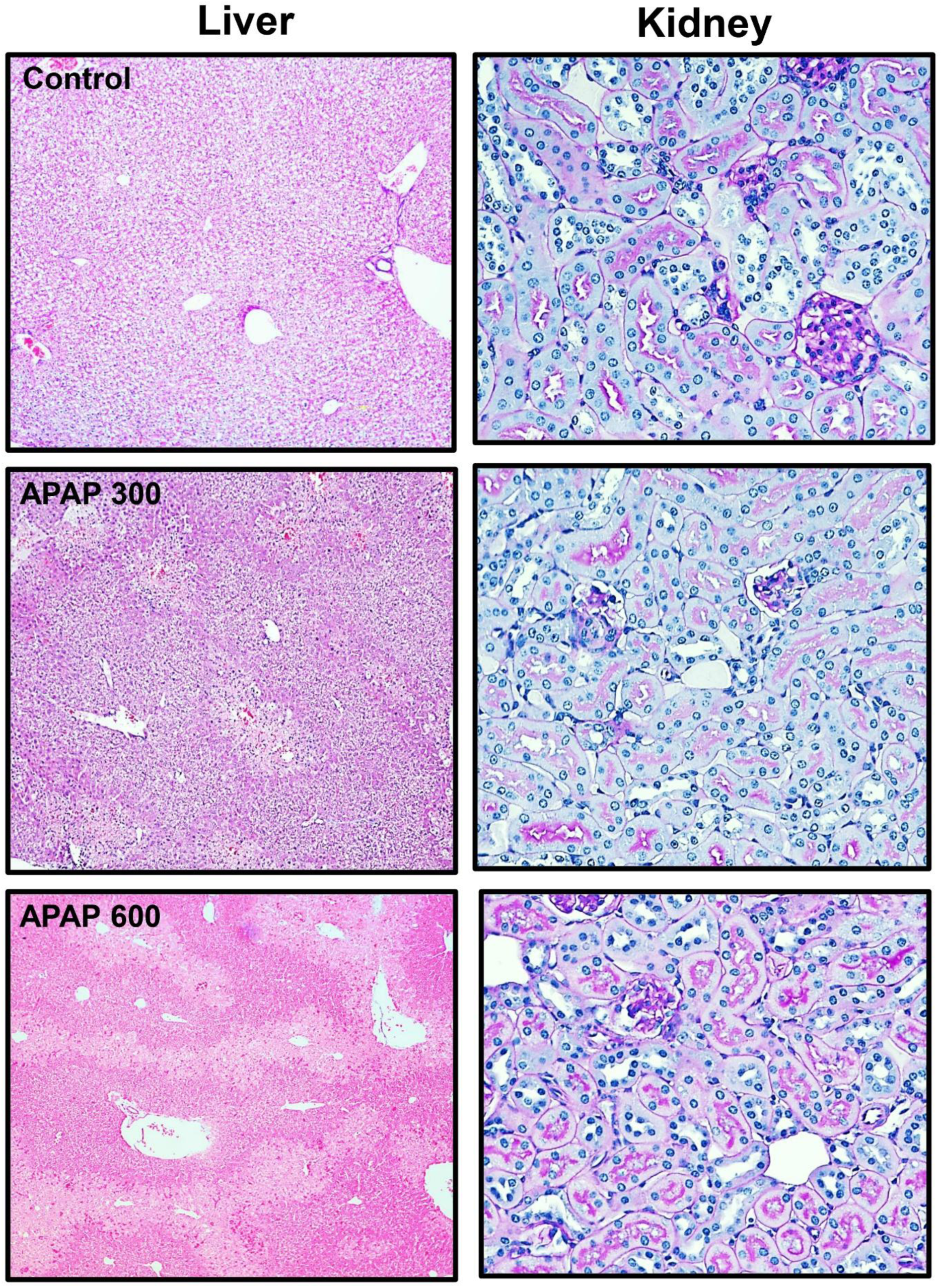
APAP induces dose-dependent hepatocellular necrosis but no detectable morphological kidney injury in female mice at 24. **h.** Representative H&E-stained liver sections and PAS-stained kidney sections from overnight-fasted female C57BL/6J mice treated with a single intraperitoneal injection of APAP (300 or 600 mg/kg) and euthanized 24 h later. Liver sections demonstrate dose-dependent centrilobular hepatocellular necrosis, which is more extensive following 600 mg/kg APAP. In contrast, PAS-stained kidney sections show preserved tubular architecture with no evidence of tubular necrosis or other overt morphological abnormalities at either APAP dose. Representative images are shown (liver, 10x; kidney, 40x).

### 3.2 GSH depletion with BSO did not amplify APAP-induced kidney injury

To investigate the contribution of glutathione depletion to APAP-induced liver and kidney injury in female mice, we used L-buthionine sulfoximine (BSO), an inhibitor of γ-glutamyl cysteine synthetase, the rate-limiting enzyme in GSH synthesis (Drew & Miners, 1984; Meister, 1988). A dose of 50 mg/kg BSO was selected based on our previous study in male mice, enabling direct comparison between sexes under identical experimental conditions (Etemadi et al., 2026a).

In overnight-fasted female mice, BSO pretreatment markedly enhanced APAP-induced hepatotoxicity. Mice receiving 300 mg/kg APAP alone developed moderate liver injury, whereas BSO pretreatment caused a dramatic increase in serum ALT activity, reaching approximately 10,000 U/L at 6 h and remaining elevated at approximately 6,000 U/L at 24 h (Figure 3A). To assess renal function, serum BUN and creatinine levels were measured. Both APAP alone and APAP+BSO treatment caused a modest increase in BUN at 3 h. However, BSO pretreatment further increased BUN levels at 6 h compared with APAP alone. This effect was transient, and BUN values returned to near-baseline levels by 24 h in both groups (Figure 3B). Serum creatinine concentrations remained unchanged throughout the study, indicating no significant renal dysfunction (Figure 3C).

**Figure 3.**
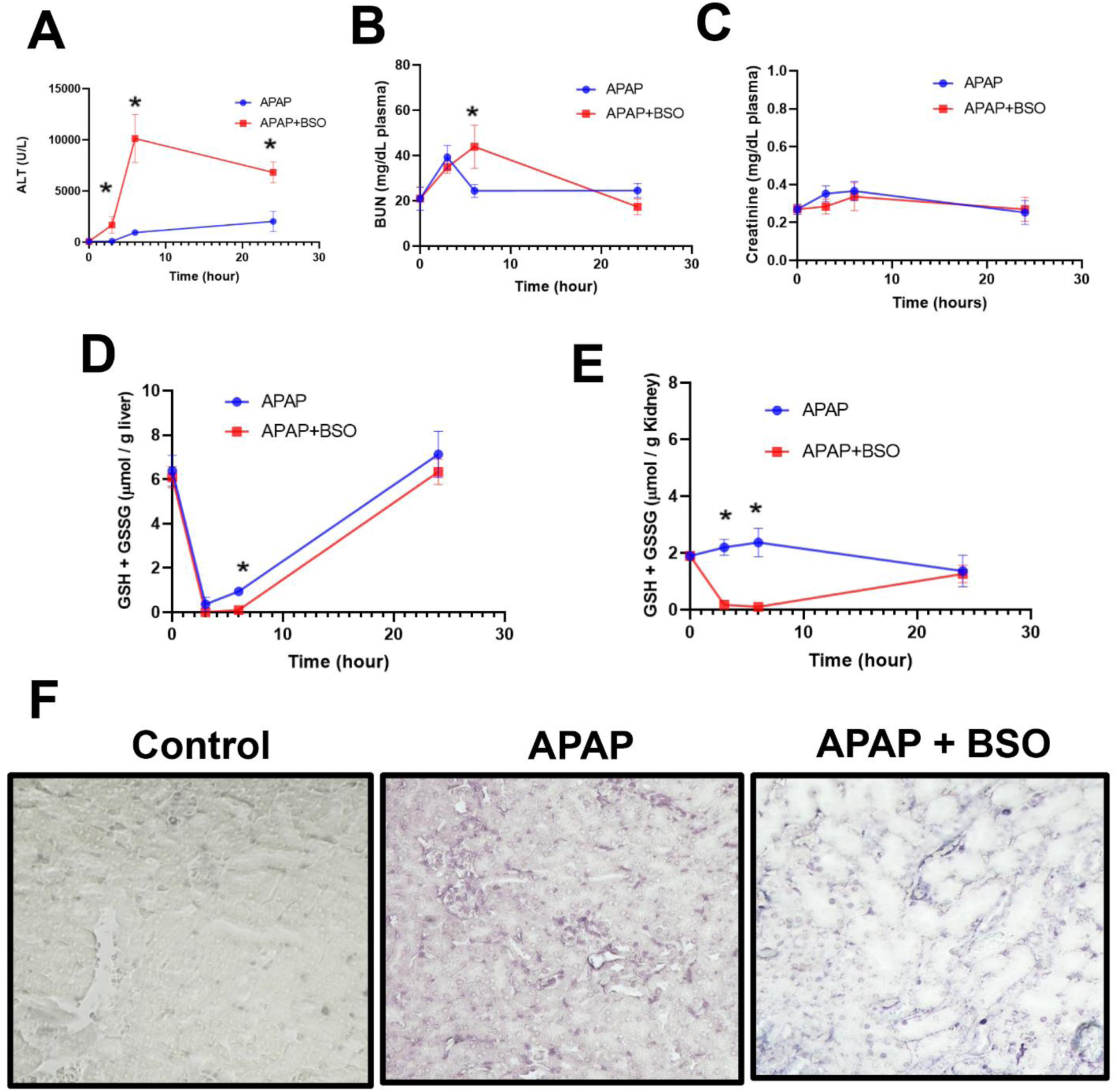
BSO exacerbates APAP-induced hepatotoxicity but does not enhance kidney injury in female mice. Overnight-fasted female C57BL/6J mice received a single intraperitoneal injection of APAP (300 mg/kg) with or without BSO pretreatment (50 mg/kg, i.p., 2 h before APAP) and were euthanized 3, 6, or 24 h after APAP administration. (A) Plasma ALT level, (B) blood urea nitrogen (BUN), (C) plasma creatinine, (D) hepatic glutathione, (E) renal glutathione, and (F) representative TUNEL-stained kidney sections at 24 h (40x). Data are presented as mean ± SEM (n = 4–5 mice/group). \**P* < 0.05 versus the APAP-treated group.

Total glutathione levels were measured in the liver and kidney. APAP alone caused a rapid decline in hepatic glutathione levels at 3 h, followed by substantial recovery by 24 h (Figure 3D). BSO pretreatment further enhanced glutathione depletion, reducing hepatic glutathione to nearly undetectable levels at both 3 and 6 h. Hepatic glutathione levels recovered substantially by 24 h.

In the kidney, APAP treatment alone had little effect on total glutathione levels at any time point. In contrast, BSO pretreatment caused near-complete depletion of renal glutathione at 3 and 6 h after APAP administration. Similar to the liver, renal glutathione levels recovered substantially by 24 h (Figure 3E). BSO alone did not affect serum ALT, BUN, creatinine, or tissue glutathione levels at 24 h (data not shown), consistent with our previous findings in male mice (Etemadi et al., 2026a).

To determine whether the profound early depletion of renal glutathione resulted in tubular cell injury, TUNEL staining was performed at 24 h. Very few TUNEL-positive cells were detected in either the APAP or APAP+BSO groups, with no appreciable difference between treatments (Figure 3F). These findings demonstrate that profound depletion of renal glutathione did not increase tubular cell death or exacerbate APAP-induced kidney injury within 24 h of treatment.

Histopathological evaluation further confirmed the differential effects of BSO pretreatment on the liver and kidney. Consistent with the serum ALT data, APAP-treated mice exhibited centrilobular hepatocellular necrosis, which was markedly exacerbated by BSO pretreatment (Figure 4). Liver sections from the BSO+APAP group showed extensive centrilobular necrosis with occasional hemorrhage, indicating more severe hepatic injury than APAP alone. In contrast, PAS-stained kidney sections revealed no apparent morphological differences between the APAP and BSO+APAP groups. Renal tubular architecture remained preserved, with no evidence of tubular necrosis or other structural abnormalities (Figure 4). To further evaluate tubular injury, kidney sections were immunostained for KIM-1 and OPN, established markers of proximal tubular injury (Cen et al., 2017; Khamissi et al., 2022; Tanase et al., 2019; Wajda et al., 2020; Xie et al., 2001). Consistent with the absence of histological injury, kidneys from both APAP- and BSO+APAP-treated mice exhibited only weak KIM-1 staining in proximal tubules and minimal OPN expression, with no apparent differences between the two groups (Figure 5). These findings indicate that BSO pretreatment did not enhance tubular injury in female mice following 300 mg/kg APAP.

**Figure 4:**
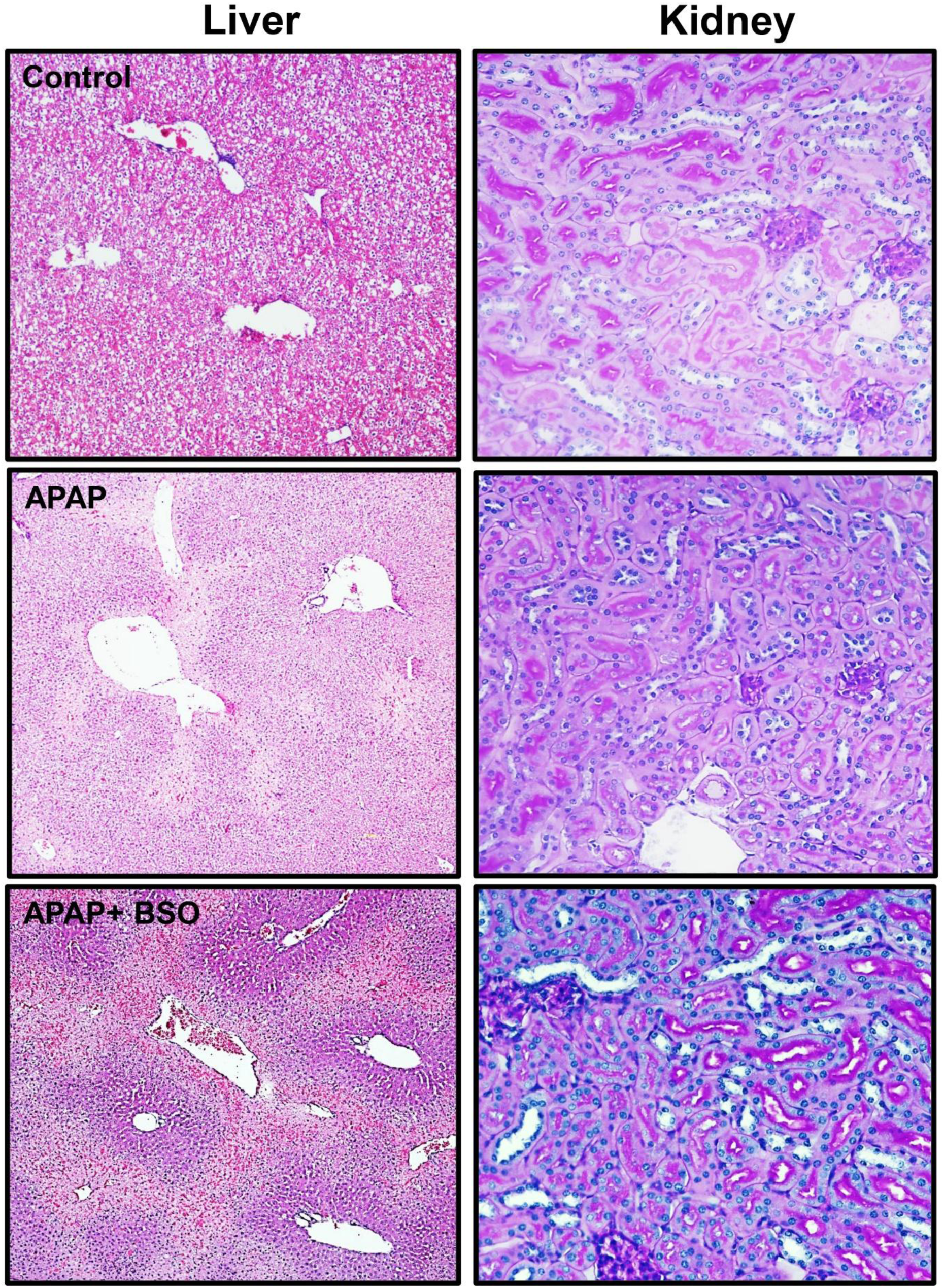
Effects of APAP and BSO on liver and renal histology. Representative H&E-stained liver sections (left) and PAS-stained kidney sections (right) from control, APAP (300 mg/kg), and APAP + BSO (50 mg/kg + 300 mg/kg APAP) female mice collected 24 h after treatment. APAP induced severe centrilobular hepatocellular necrosis, which was further exacerbated by BSO pretreatment. In contrast, PAS-stained kidney sections show preserved tubular architecture with no evidence of tubular necrosis or other overt morphological abnormalities at either APAP dose. Representative images are shown (liver, 10x; kidney, 40x).

**Figure 5.**
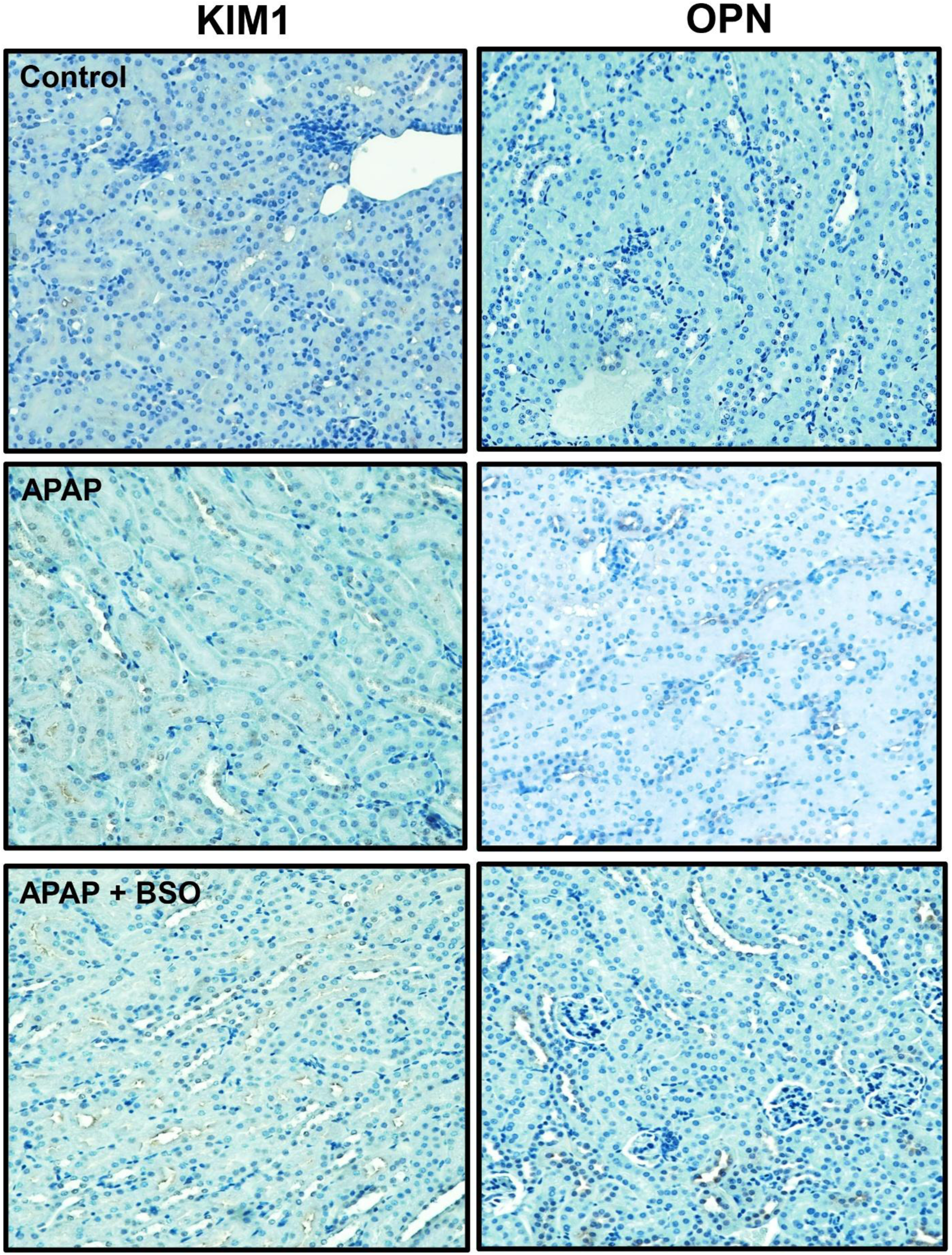
Representative immunohistochemical staining of KIM-1 and OPN 24 h after APAP treatment. Representative kidney sections immunostained for KIM-1 (left, 40x) and OPN (right, 40x) from control, APAP (300 mg/kg), and APAP + BSO (50 mg/kg + 300 mg/kg APAP) female mice collected 24 h after treatment. Minimal KIM-1 and OPN immunoreactivity was observed in all groups, consistent with the absence of substantial renal tubular injury. Images are representative of n = 4–5 mice per group.

### 3.3. Delayed progression of APAP-induced acute kidney injury prevented by 4-MP but not NAC

APAP-induced AKI frequently develops and progresses beyond the initial 24 h after overdose (Jones & Vale, 1993; Mour et al., 2005). However, severe hepatotoxicity in the mouse model limits survival at these later time points (Akakpo et al., 2023; Bhushan et al., 2014; Nguyen et al., 2023), precluding assessment of delayed nephrotoxicity. Therefore, mice were treated with NAC, which improves survival but has limited efficacy against APAP-induced early AKI (Etemadi et al., 2026a), to enable evaluation of delayed kidney injury at 48 h. In parallel, female mice were co-treated with 4-MP to determine whether 4-MP remains protective against APAP-induced kidney injury in the absence of detectable renal CYP2E1 expression.

Female C57BL/6J mice received 600 mg/kg APAP with or without NAC (500 mg/kg), and liver and kidney injury were assessed at 24 and 48 h after treatment (Figure 6). Consistent with its known hepatoprotective effect, NAC markedly attenuated APAP-induced hepatotoxicity, reducing serum ALT activities by approximately 75-80% at both 24 and 48 h compared with APAP alone (Figure 6A). Despite the marked attenuation of hepatotoxicity by NAC, renal dysfunction continued to progress. Serum BUN remained elevated in the APAP+NAC group at both 24 and 48 h, with the highest levels observed at 48 h (Figure 6B). Interestingly, BUN was elevated with NAC alone at both 24 and 48 h. Likewise, serum creatinine progressively increased over time and reached its highest level in the APAP+NAC group at 48 h (Figure 6C). Creatinine levels were also significantly elevated with NAC treatment alone, both at 24 and 48 h. However, histopathological examination of PAS-stained kidney sections from NAC-alone-treated mice revealed preserved tubular architecture without apparent tubular vacuolation or dilation at either time point (Supplementary Figure 1). Immunohistochemical analysis showed modest increases in KIM-1 and OPN expression following NAC treatment alone, whereas TUNEL staining revealed minimal tubular cell death (Supplementary Figures 1 and 2). These findings suggest that the elevations in renal functional markers following NAC administration were not accompanied by substantial structural kidney injury. Kidney sections revealed minimal morphological alterations in APAP-treated mice at 24 h following NAC administration, with renal architecture largely comparable to control animals (Figure 6D). In contrast, by 48 h, kidneys from the APAP+NAC-treated group exhibited clear evidence of tubular injury, characterized by tubular vacuolation (black arrows), consistent with the progressive increases in serum BUN and creatinine (Figure 6D). Consistent with the histopathological findings, KIM-1, OPN, and TUNEL staining showed minimal changes at 24 h following APAP+NAC treatment (Figure 7).

**Figure 6.**
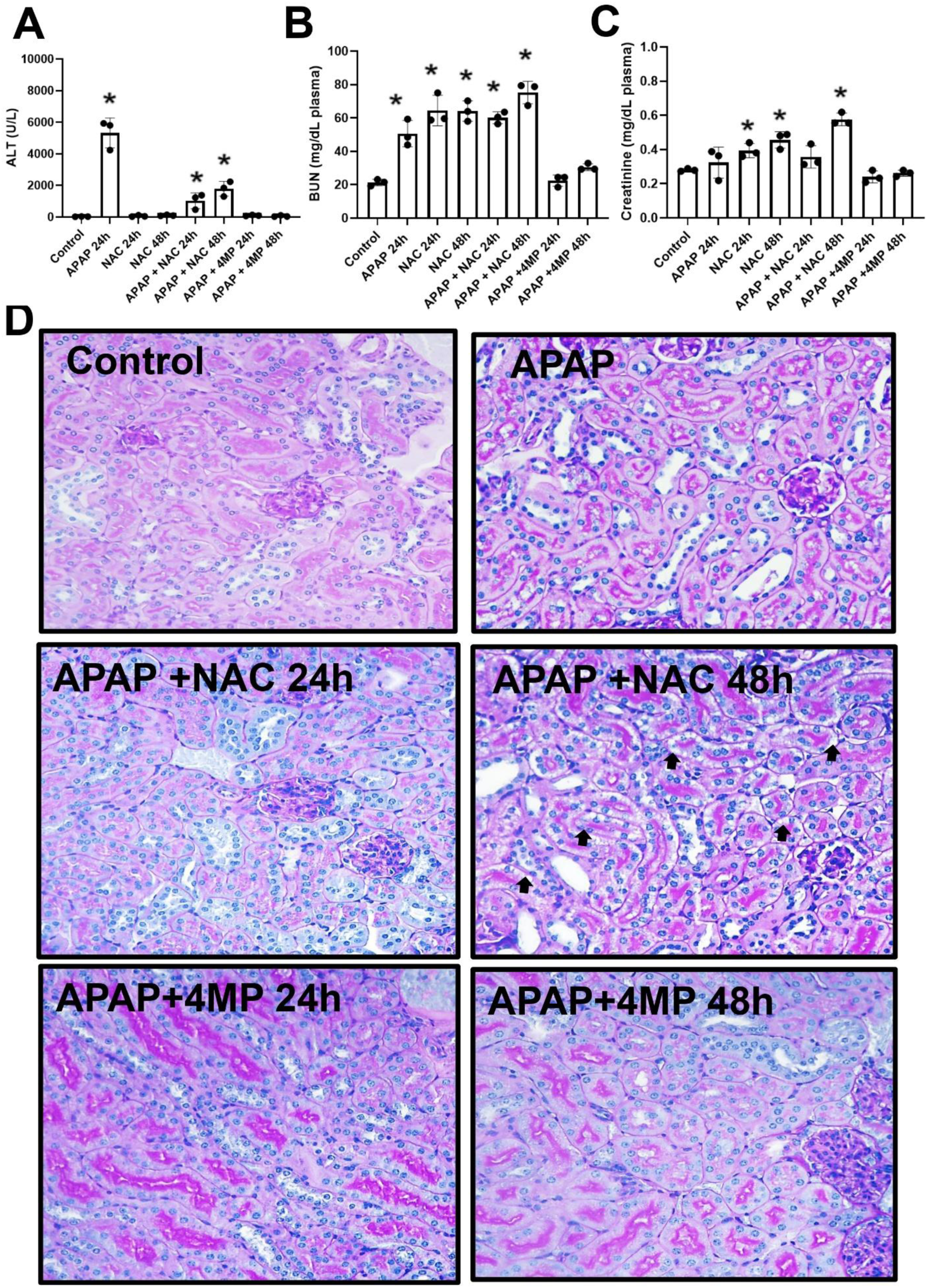
Fomepizole (4-MP), but not NAC, prevents delayed APAP-induced kidney injury in female mice. Overnight-fasted female C57BL/6J mice received APAP (600 mg/kg, i.p.) alone or in combination with NAC (500 mg/kg, i.p.) or 4-MP (50 mg/kg, i.p.) and were evaluated at 24 or 48 h after APAP administration. (A) Plasma ALT activity, (B) blood urea nitrogen (BUN), (C) plasma creatinine, and (D) representative PAS-stained kidney sections. Arrows indicate tubular vacuolation in the APAP+NAC group at 48 h. Data are presented as mean ± SEM (n = 4 mice/group). *P < 0.05 versus control. Representative histological images are shown at 40× magnification.

**Figure 7.**
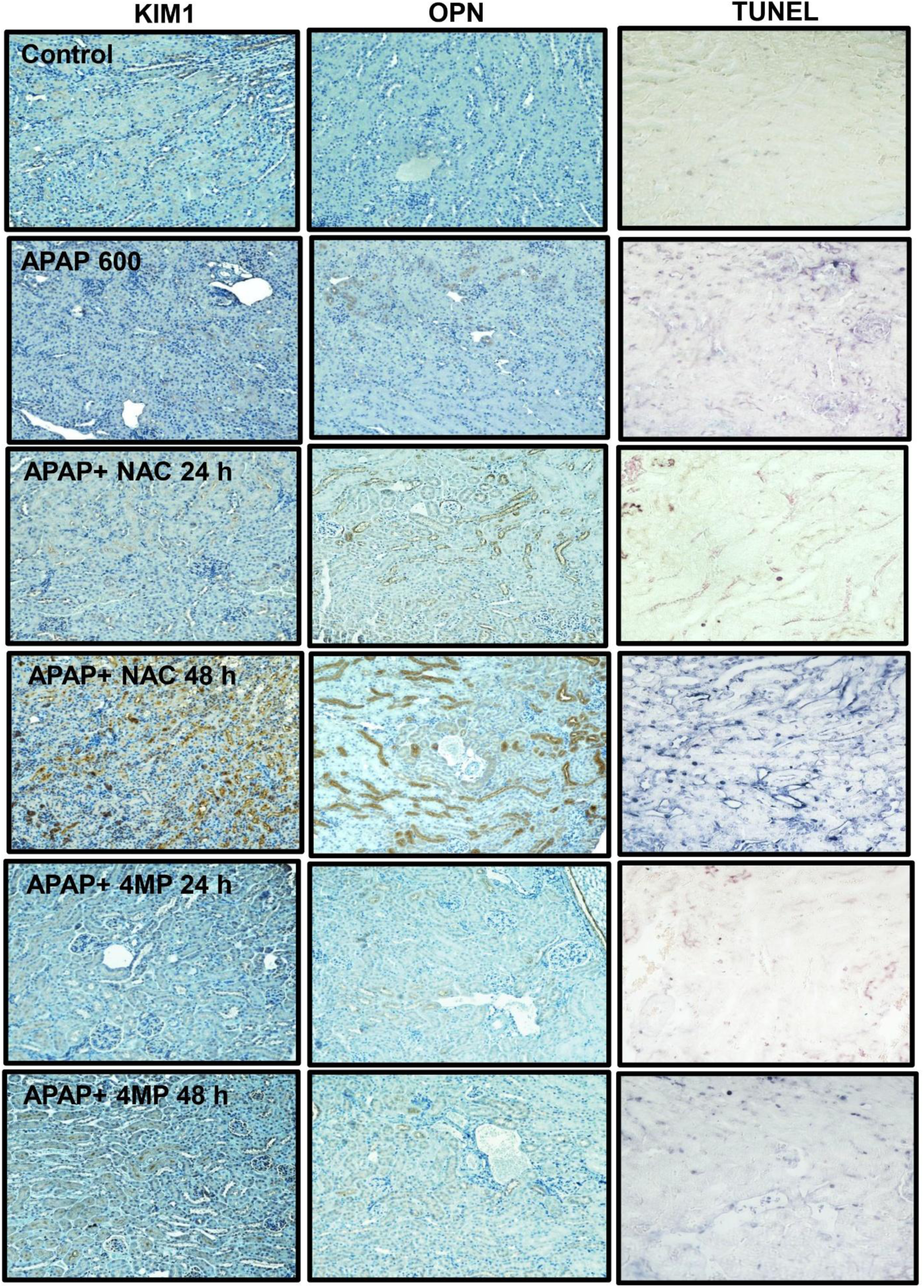
Fomepizole (4-MP) prevents delayed tubular injury and cell death following APAP overdose in female mice. Representative kidney sections immunostained for KIM-1 (left) and OPN (middle), and TUNEL staining (right) from control, APAP (600 mg/kg), APAP+NAC, and APAP+4-MP groups. APAP+NAC treatment was associated with increased tubular injury at 48 h, whereas KIM-1 and OPN expression and TUNEL-positive cells remained minimal following 4-MP treatment. Representative images are shown at 20× for KIM-1 and OPN and 40× for TUNEL (n = 4 mice/group).

However, by 48 h, KIM-1 and OPN expression were markedly increased and accompanied by numerous TUNEL-positive tubular cells, indicating substantial tubular injury and cell death (Figure 7). Conversely, 4-MP provided sustained protection against both hepatic and renal injury (Figure 6,7). Serum ALT activities remained near control levels at both 24 and 48 h in APAP+4-MP-treated mice (Figure 6A). Importantly, 4-MP also prevented the progressive increase in renal dysfunction, with BUN and creatinine remaining near control levels at both time points (Figure 6B,C). Consistent with these functional measurements, kidney morphology was largely preserved following 4-MP treatment at both 24 and 48 h, with substantially less tubular injury compared with the APAP+NAC group at 48 h (Figure 6D). KIM-1 and OPN staining also remained minimal following 4-MP treatment, and few TUNEL-positive tubular cells were observed at either time point (Figure 7). Thus, 4-MP prevented the delayed progression of APAP-induced AKI even in female mice lacking detectable renal CYP2E1 expression. Together, these results demonstrate that delayed APAP-induced nephrotoxicity develops despite NAC-mediated hepatoprotection, whereas 4-MP prevents the progression of renal injury even in the absence of detectable renal CYP2E1 expression.

Finally, histopathological scoring of the renal cortex and outer stripe of the outer medulla (OSOM) was performed by a renal expert pathologist (T.A.F.) to assess the severity of tubular injury. Most treatment groups exhibited minimal histopathological injury (scores 0–1). However, at 48 h following APAP+NAC treatment, three of four mice received a score of 2, consistent with the increased tubular injury observed by PAS staining (Supplementary Figure 3).

## 4. Discussion

### 4.1 Female mice are resistant to APAP-induced AKI despite severe hepatotoxicity at 24 h

APAP-induced AKI is a relatively delayed complication that is most commonly observed in patients with severe hepatotoxicity and acute liver failure, a pattern that is well reproduced in experimental mouse models (Akakpo et al., 2020; Hart et al., 1994, 1995). This close temporal association has led to the hypothesis that severe liver injury contributes to the development or progression of renal injury. However, clinical reports describing APAP-induced AKI in the absence of acute liver failure indicate that nephrotoxicity can also occur independently of severe hepatotoxicity (Eguia & Materson, 1997; Hart et al., 1994, 1995). Consistent with this concept, previous mechanistic studies, including our own, have demonstrated that the kidney can locally bioactivate APAP through CYP2E1, resulting in NAPQI formation, protein adduct formation, endoplasmic reticulum stress, and apoptotic death of proximal tubular cells (Akakpo et al., 2020, 2023; Etemadi et al., 2026a). To distinguish the contribution of severe liver injury from that of local renal bioactivation, we used female mice, which naturally lack CYP2E1 expression in proximal tubular cells (Hoivik et al., 1995; Hu et al., 1993) and are therefore largely protected from early CYP2E1-dependent nephrotoxicity after APAP overdose (Etemadi et al., 2026b). This unique model allowed us to determine whether severe hepatotoxicity alone is sufficient to induce AKI in the absence of renal CYP2E1-mediated bioactivation. Consistent with our previous findings in male mice, female mice developed severe hepatotoxicity but exhibited only minimal kidney injury 24 h after APAP overdose (Figures 1, 2), indicating that hepatotoxicity alone is insufficient to induce early AKI and that local renal bioactivation plays a critical role.

### 4.2 Glutathione depletion exacerbates hepatotoxicity but does not promote AKI

Female mice are relatively resistant to APAP-induced hepatotoxicity after a moderate overdose because they recover hepatic GSH more rapidly than male mice, a response associated with enhanced induction of GCLC (Du et al., 2014). To determine whether increasing the severity of hepatotoxicity promotes AKI, we pretreated mice with BSO to inhibit GSH synthesis. Because high BSO doses markedly enhance APAP-induced hepatotoxicity and cause early mortality in female mice (Drew & Miners, 1984; Griffith & Meister, 1979; Masubuchi et al., 2011), we selected a lower dose (50 mg/kg) that we previously demonstrated produces substantial early hepatic GSH depletion while preserving survival following APAP overdose (Etemadi et al., 2026a). As expected, BSO accelerated hepatic and renal GSH depletion during the first 6 h after APAP administration, with GSH levels recovering by 24 h. Despite exacerbating hepatotoxicity, BSO did not increase renal dysfunction or kidney injury at 24 h (Figure 3). These findings indicate that increasing the severity of hepatotoxicity alone is insufficient to trigger early AKI, further supporting the conclusion that APAP-induced liver and kidney injury are mechanistically distinct processes. In contrast to our previous findings in male mice (Etemadi et al., 2026a), profound early depletion of renal GSH was insufficient to promote kidney injury in female mice. This observation is consistent with the absence of renal CYP2E1 expression and the lack of detectable renal APAP-protein adduct formation during the early phase after APAP overdose (Etemadi et al., 2026b).

Together, these findings indicate that local CYP2E1-dependent NAPQI formation, rather than hepatotoxicity or glutathione depletion alone, is the primary determinant of early APAP-induced nephrotoxicity in male but not in female mice.

### 4.3 Delayed kidney injury develops despite NAC-mediated hepatoprotection

To determine whether severe hepatotoxicity is required for the progression of APAP-induced AKI, we administered NAC following a 600 mg/kg APAP overdose. Because severe liver injury markedly limits survival beyond 24 h, NAC treatment enabled evaluation of delayed kidney injury at later time points. Using this approach, we found that substantial AKI developed by 48 h despite marked protection against liver injury. In parallel, we evaluated 4-MP to determine whether it could prevent delayed kidney injury in female mice despite the absence of detectable renal CYP2E1 expression.

NAC is the standard therapy for APAP overdose and exerts its hepatoprotective effects primarily through replenishment of glutathione stores, enhancement of NAPQI detoxification, and scavenging of mitochondrial oxidant stress (Corcoran & Wong, 1986; Saito et al., 2010). Consistent with these established mechanisms, NAC markedly reduced serum ALT activities and improved survival in APAP-treated female mice. However, despite this robust hepatoprotection, substantial kidney injury became evident at 48 h, as indicated by elevated serum BUN and creatinine, tubular vacuolation, increased KIM-1 and OPN expression, and extensive tubular cell death (Figures 6, 7). It should also be noted that high-dose NAC alone produced a modest increase in BUN and creatinine, consistent with our previous findings (Etemadi et al., 2026a) and indicating that mild renal functional changes following NAC administration are not unprecedented. Similar observations have been reported with high experimental doses of NAC, which have been associated with tubular injury, increased oxidant stress, and impaired glutathione homeostasis (Tsai et al., 2024). In addition, excessive antioxidant supplementation may interfere with adaptive Nrf2-mediated stress responses, potentially limiting endogenous cytoprotective mechanisms (Small et al., 2018). Consistent with these observations, NAC treatment alone in our study induced modest increases in the tubular injury markers KIM-1 and OPN without evidence of TUNEL-positive cells or tubular vacuolation on PAS-stained kidney sections (Supplementary Figures 1, 2), suggesting a mild sublethal tubular stress response rather than overt tubular injury or cell death.

The delayed appearance of AKI in NAC-treated mice closely reproduces the temporal pattern observed in patients after APAP overdose, in whom renal dysfunction frequently becomes evident several days after the onset of hepatotoxicity (Jones & Vale, 1993; Mazer & Perrone, 2008; Mour et al., 2005). Because NAC remains the standard treatment for APAP overdose but provides limited protection against AKI, our findings underscore the clinical relevance of this model for investigating the mechanisms responsible for delayed nephrotoxicity that are not prevented by current therapy. Importantly, the failure of NAC to prevent delayed AKI despite robust hepatoprotection indicates that the mechanisms driving delayed nephrotoxicity are fundamentally distinct from those responsible for the early phase of injury. Whereas early APAP-induced nephrotoxicity depends on local renal CYP2E1-mediated NAPQI formation, the delayed injury observed in female mice likely reflects CYP2E1-independent pathways.

4-MP or fomepizole is a competitive alcohol dehydrogenase inhibitor clinically used for methanol and ethylene glycol poisoning (McMartin et al., 2025) that also inhibits CYP2E1-dependent APAP bioactivation (Akakpo et al., 2018). In APAP hepatotoxicity, 4-MP prevents NAPQI formation when administered early and can also protect after the metabolic phase by inhibiting JNK activation and its mitochondrial translocation (Akakpo et al., 2019). 4-MP also protects against APAP-induced nephrotoxicity in male mice by inhibiting renal CYP2E1-dependent APAP bioactivation and subsequent ER stress-mediated apoptosis (Akakpo et al., 2020, 2023). Importantly, unlike the liver, APAP-induced kidney injury is not characterized by substantial mitochondrial protein adduct formation or JNK activation, indicating distinct downstream mechanisms (Akakpo et al., 2020, 2023). In the present study, 4-MP prevented renal injury at both 24 and 48 h despite the absence of detectable renal CYP2E1 expression in female mice. This finding suggests that inhibition of renal CYP2E1 alone cannot explain the renoprotective effect of 4-MP and raises the possibility of an additional CYP2E1-independent mechanism. Because the JNK-dependent mitochondrial pathway targeted by 4-MP in the liver does not appear to operate similarly in the kidney, the mechanism responsible for this protection remains unclear and may involve an alternative metabolic or downstream cellular stress pathway.

Although our findings support a transition from early CYP2E1-dependent to delayed CYP2E1-independent nephrotoxicity, the mechanisms responsible for the latter are likely multifactorial. APAP can undergo deacetylation by renal N-deacetylases to form *p*-aminophenol (PAP), a well-established nephrotoxic metabolite that undergoes further bioactivation and induces oxidative stress and tubular injury independently of renal CYP2E1 (Newton et al., 1982, 1985). Consistent with this mechanism, our recent findings identify PAP formation as a partial contributor to delayed APAP-induced AKI through a CYP2E1-independent pathway (Etemadi et al., 2026b). However, additional mechanisms are also likely involved. For example, prostaglandin endoperoxidase synthetase (PGES), which is highly expressed in the renal medulla, can bioactivate APAP to reactive intermediates in regions where CYP450 activity is low, thereby providing an alternative CYP2E1-independent pathway for oxidative injury (Mazer & Perrone, 2008; Sciskalska et al., 2015). Whether 4-MP interferes with these alternative metabolic pathways or acts further downstream to limit the cellular consequences of delayed APAP-induced renal injury remains to be determined. Future studies should define the relative contribution of these mechanisms to delayed APAP-induced AKI and determine whether targeting these pathways can improve renal protection.

## 5. Summary and conclusions

Our study demonstrates that APAP-induced liver injury and kidney injury are mechanistically distinct and differentially regulated in female mice. Despite severe hepatotoxicity, female mice lacking renal CYP2E1 developed minimal kidney injury during the first 24 h after APAP overdose, indicating that hepatotoxicity alone is insufficient to initiate early AKI. Likewise, pharmacological depletion of glutathione with BSO markedly exacerbated hepatotoxicity but did not promote renal injury, further supporting the critical role of local renal CYP2E1-independent bioactivation during the early phase of nephrotoxicity. In contrast, NAC substantially attenuated liver injury and improved survival but failed to prevent delayed AKI, demonstrating that kidney injury continues to progress despite effective hepatoprotection. Notably, 4-MP prevented renal injury at both 24 and 48 h despite the absence of detectable renal CYP2E1 expression in female mice. This finding suggests that the renoprotective effects of 4-MP may extend beyond inhibition of CYP2E1-mediated APAP bioactivation and involve additional CYP2E1-independent mechanisms. Collectively, these findings support a temporal model in which early APAP-induced AKI in male mice depends on renal CYP2E1-mediated NAPQI formation, whereas delayed nephrotoxicity in both male and female animals is driven by CYP2E1-independent mechanisms, including PAP formation and potentially other alternative bioactivation pathways. These findings highlight the need for therapeutic strategies that directly target renal injury mechanisms in addition to protecting the liver following APAP overdose.

## Acknowledgements

This work was funded in part by the National Institute of Diabetes and Digestive and Kidney Diseases (NIDDK) grants R01 DK102142 and DK125465, and National Institute of General Medical Sciences (NIGMS) funded Liver Disease COBRE grants P20 GM103549 and P30 GM118247. Y.E. was supported in part by the Biomedical Research Training Program (BRTP) at the University of Kansas Medical Center.

## Credit authorship contribution statement

Yasaman Etemadi: Writing – original draft, Methodology, Investigation, Data curation. Timothy A. Fields: Writing – review & editing, Investigation, Formal analysis. Anup Ramachandran: Writing – review & editing, Visualization, Validation, Investigation, Funding acquisition, Data curation. Hartmut Jaeschke: Writing – review & editing, Validation, Supervision, Project administration, Funding acquisition, Conceptualization.

## Supplemental Figure Legends

**Figure S1:**
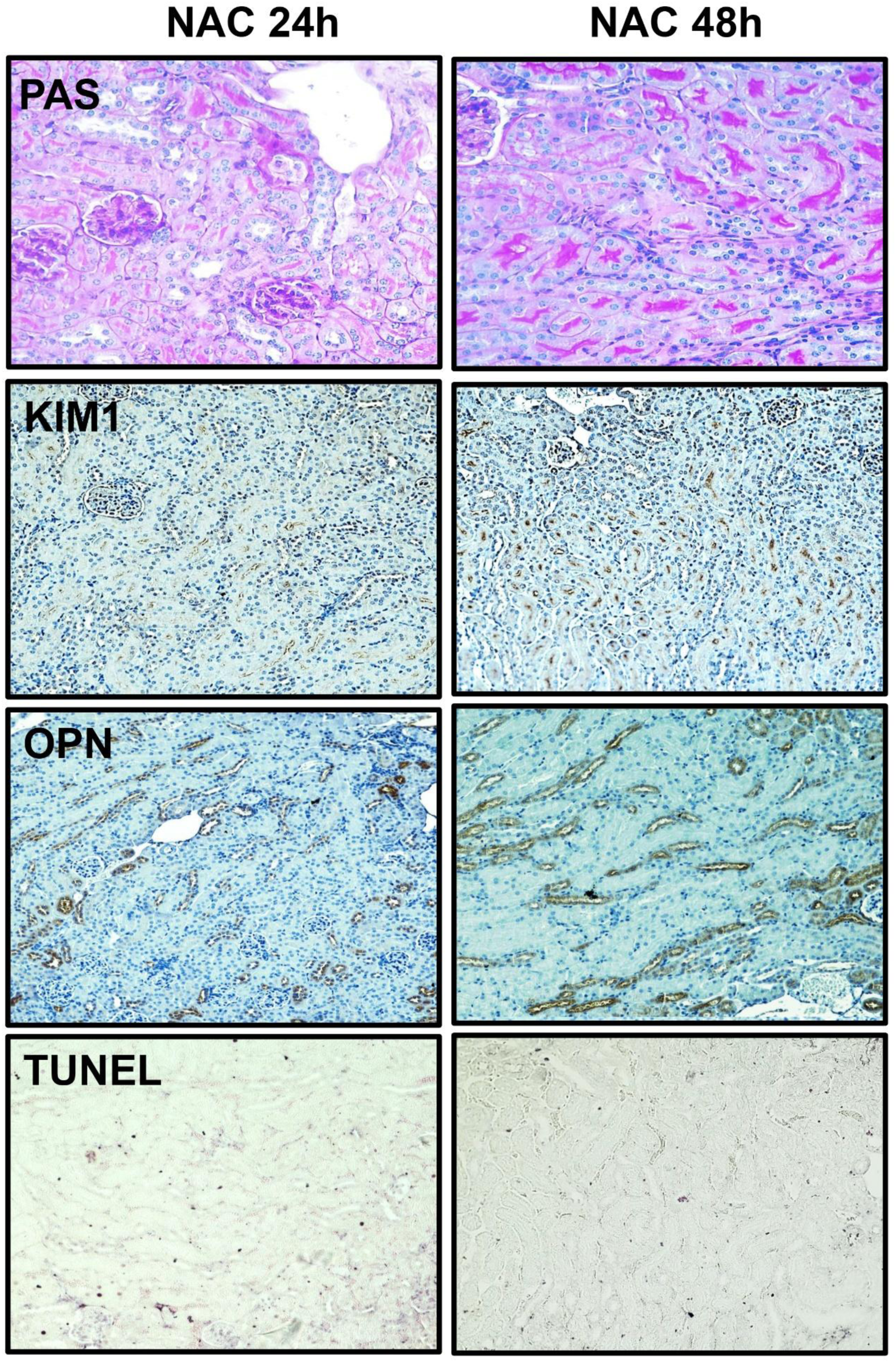
Histopathological assessment of renal injury following NAC treatment in female mice. Female C57BL/6J mice received NAC (500 mg/kg) and were euthanized at 24 or 48 h. We evaluated representative kidney sections by PAS staining to assess tubular morphology, immunohistochemistry for KIM-1 and OPN to assess tubular injury, and TUNEL staining to assess tubular cell death.

**Figure S2:**
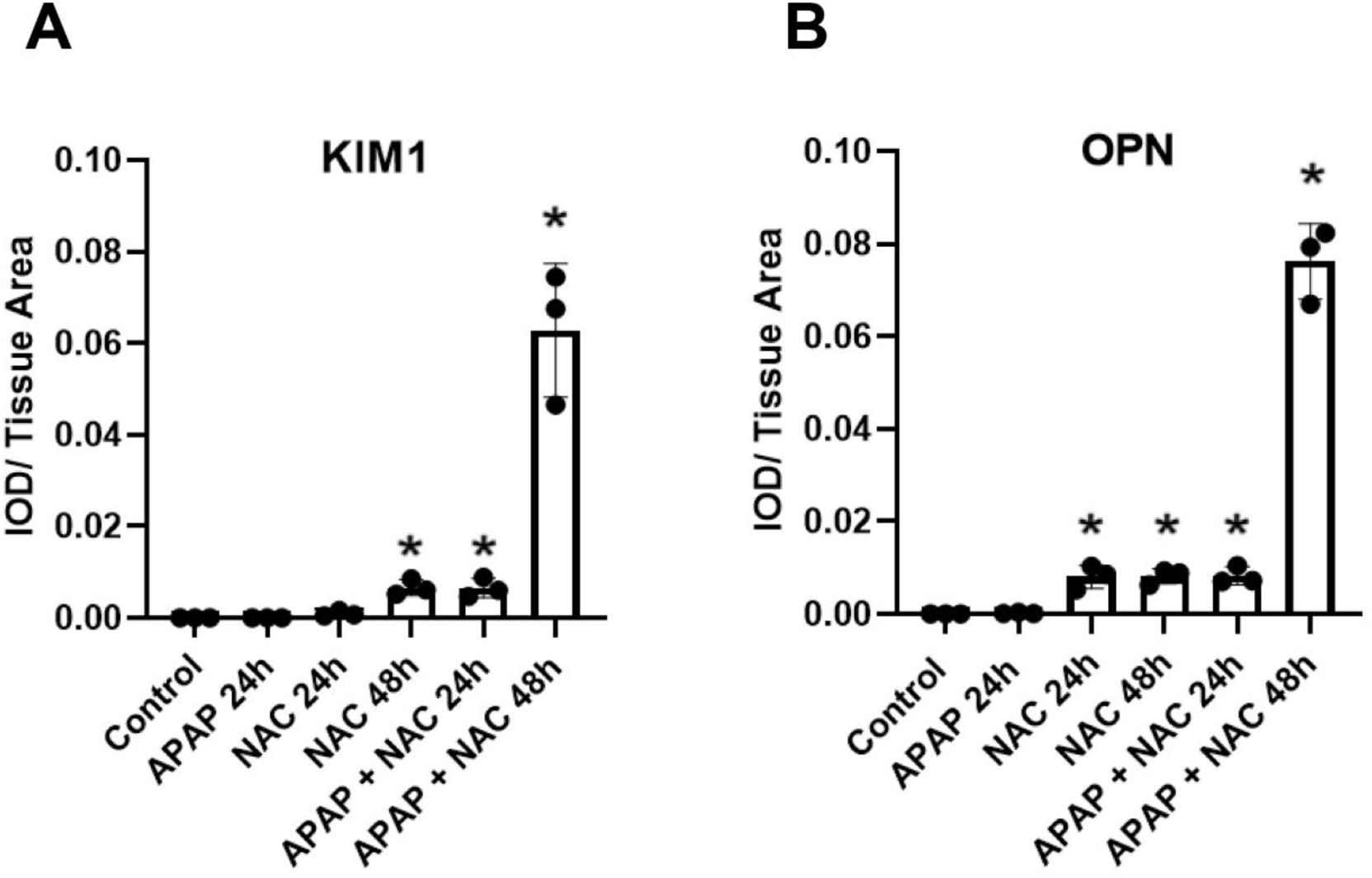
Quantification of renal KIM-1 and osteopontin immunostaining following APAP and NAC treatment in female mice. Female C57BL/6J mice received APAP (600 mg/kg), NAC (500 mg/kg), or APAP + NAC and were euthanized at 24 or 48 h after treatment. Immunohistochemical staining for (A) KIM-1 and (B) OPN was quantified using Fiji (ImageJ). We normalized integrated optical density (IOD) to total tissue area (IOD/tissue area). For each mouse, 3-4 non-overlapping fields of the renal cortex were analyzed at 20× magnification. Data are presented as mean ± SEM (n = 3-4 mice/group). *p < 0.05 compared with controls.

**Figure S3:**
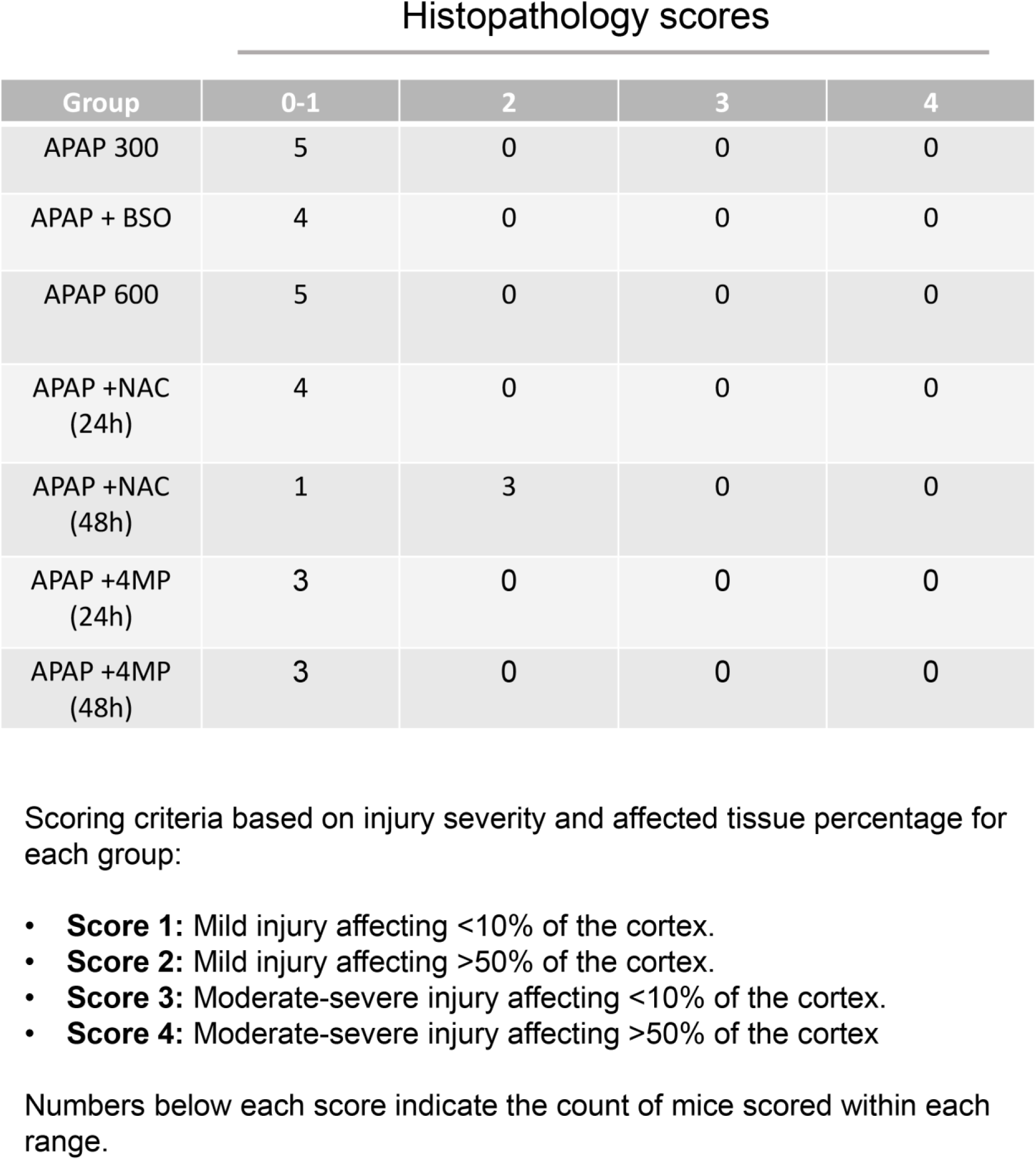
Histopathological scoring of renal injury in female mice following APAP, APAP + BSO, APAP + NAC, and APAP + 4-MP treatment. We assessed renal injury based on the severity and extent of tubular damage. Scores ranged from 0-1 (minimal injury) to 4 (severe injury). Values represent the number of mice assigned to each score within each treatment group.

